# Survey of human chromosome 21 gene expression effects on early development in *Danio rerio*

**DOI:** 10.1101/269944

**Authors:** Sarah McGuire Edie, Norann A Zaghloul, Carmen C Leitch, Donna K Klinedinst, Janette Lebron, Joey F Thole, Andrew S McCallion, Nicholas Katsanis, Roger H. Reeves

**Affiliations:** Johns Hopkins University School of Medicine, Institute of Genetic Medicine; University of Maryland School of Medicine, Department of Medicine, Division of Endocrinology, Diabetes and Nutrition; Johns Hopkins University School of Medicine, Department of Physiology; Center for Human Disease Modeling, Duke University

## Abstract

Trisomy for human chromosome 21 (Hsa21) results in Down syndrome (DS), one of the most genetically complex conditions compatible with human survival. Assessment of the physiological consequences of dosage-driven overexpression of individual Hsa21 genes during early embryogenesis and the resulting contributions to DS pathology in mammals are not tractable in a systematic way. A recent study looked loss-of-function of *C. elegans* orthologues of Hsa21 genes and identified ten candidates with behavioral phenotypes, but the equivalent over-expression experiment has not been done. We turned to zebrafish as a developmental model and, using a number of surrogate phenotypes, we screened Hsa21 genes for dosage sensitive effects on early embyrogenesis. We prepared a library of 164 cDNAs of conserved protein coding genes, injected mRNA into early embryos and evaluated up to 5 days post-fertilization (dpf). Twenty-four genes produced a gross morphological phenotype, 11 of which could be reproduced reliably. Seven of these gave a phenotype consistent with down regulation of the sonic hedgehog (Shh) pathway; two showed defects indicative of defective neural crest migration; one resulted consistently in pericardial edema; and one was embryonic lethal. Combinatorial injections of multiple Hsa21 genes revealed both additive and compensatory effects, supporting the notion that complex genetic relationships underlie end phenotypes of trisomy that produce DS. Together, our data suggest that this system is useful in the genetic dissection of dosage-sensitive gene effects on early development and can inform the contribution of both individual loci and their combinatorial effects to phenotypes relevant to the etiopathology of DS.

## INTRODUCTION

Down syndrome (DS) occurs in about one of 700 live births due to trisomy for human chromosome 21 (Hsa21) [1]. The consequent ~1.5 fold over expression of most genes on Hsa21 can result in more than 80 clinical phenotypes, many of which originate during prenatal development and vary in both severity and penetrance [2–4]. Among the most consistent features are cognitive impairment, characteristic craniofacial dysmorphism, smaller and hypocellular brain and Alzheimer histopathology [5, 6]. Individuals with DS also have a greatly increased risk of congenital heart disease, Hirschsprung disease and acute megakaryoblastic leukemia in children. However, the incomplete penetrance of many DS phenotypes indicates that trisomy 21 is not sufficient to cause most of these conditions, suggesting an important role for allelic variation of Hsa21 genes and additional modifier genes, as well as potential environmental and stochastic factors [7–9]. DS is one of the few autosomal aneuploidies that is compatible with life, likely related to Hsa21 being the smallest autosome and having very low gene density. Estimates of the gene content on Hsa21 range from ~300-600 genes/transcripts, of which 162 have been identified as well-conserved in other mammals [10]. Understanding how trisomy for these genes affects the presentation of the phenotypes in DS is a major focus for research into this condition.

A major challenge in understanding mechanisms of gene action in DS is that trisomy 21 is present from conception and every cell is affected, causing effects throughout development. Trisomic genes may have a primary effect directly on cellular function or may secondarily affect expression and regulation of disomic genes. Trisomy-induced changes in one cell type could alter interactions with neighboring cells, thus initiating cascades of primary and secondary effects [6, 11]. Use of mouse models trisomic for different segments of Hsa21-orthologous sequences supports to an extent the idea that different genetic segments correlate with some specific phenotypes [12, 13] [14], but even the smallest segmental trisomy still contains many genes. The effort and cost to systematically engineer individual transgenic mouse models of all conserved genes on Hsa21 would be prohibitive, to say nothing of the analysis of the possible combinations of genes. Further, events early in embryogenesis are difficult to access in mammals.

Recently, a screen to examine the effects of down-regulating orthologs of 47 Hsa21 was performed in *Caenorhabditis elegans* [15]. Ten of these conserved genes exhibited neurobehavioral phenotypes: *COL18A1*, *CBS*, *DONSON*, *EVA1C*, *N6AMT*, *NCAM2*, *POFUT2*, *PDXK*, *RUNX1 and SYNJ1* [15]. Of these ten genes, five were shown to be essential for development based on the lethality phenotype seen in mouse knock-out models. The C. elegans screen identified three genes that were previously uncharacterized (*DONSON*, *N6AMT and PDXK*) as having a phenotype, providing new information about DS related genes and showing that these types of expression-based screens can provide a valuable resource to the DS research community. The knockdown screen in worms for all of the likely Hsa21 orthologs provided insights into gene function, but an over-expression screen that might be more relevant to over-expression in DS has not been done. Previous studies have shown that the expression and/or suppression in zebrafish embryos of genes that map to disease-associated duplications and deletions in people can distinguish potent drivers of pathology [16–20]. Motivated by such studies, we systematically over-expressed in zebrafish embryos each of 164 Hsa21 cDNAs representing 163 genes and assessed their effects on early development.

## RESULTS

### Development of the clone set and initial screen

In a detailed annotation of transcripts from Hsa21, Sturgeon and Gardiner identified 162 genes that are highly conserved with mouse [10]. We assembled a set of Hsa21 cDNA clones consisting of 148 of these conserved genes and 15 human-specific genes [10, 21]. The 15 genes that are not conserved include DSCR genes, long intergenic non-coding RNA, and Hsa21 open reading frames. One gene, *SYNJ1*, is represented by two splice isoforms, for a total of 164 cDNAs (Supplemental Table 1). The majority of clones are from the InVitrogen UltimateORF collection library (now Thermo Fisher Scientific’s Ultimate ORF Clones), subcloned into the pCS2+ destination vector (Figure 1) [22]. Thus we selected highly conserved genes from this carefully annotated set for which cDNA could be obtained and which could be cloned into the pCS2+ vector and transcribed. Additional sources and vectors are described in Supplemental Table 1.

**Figure 1:**
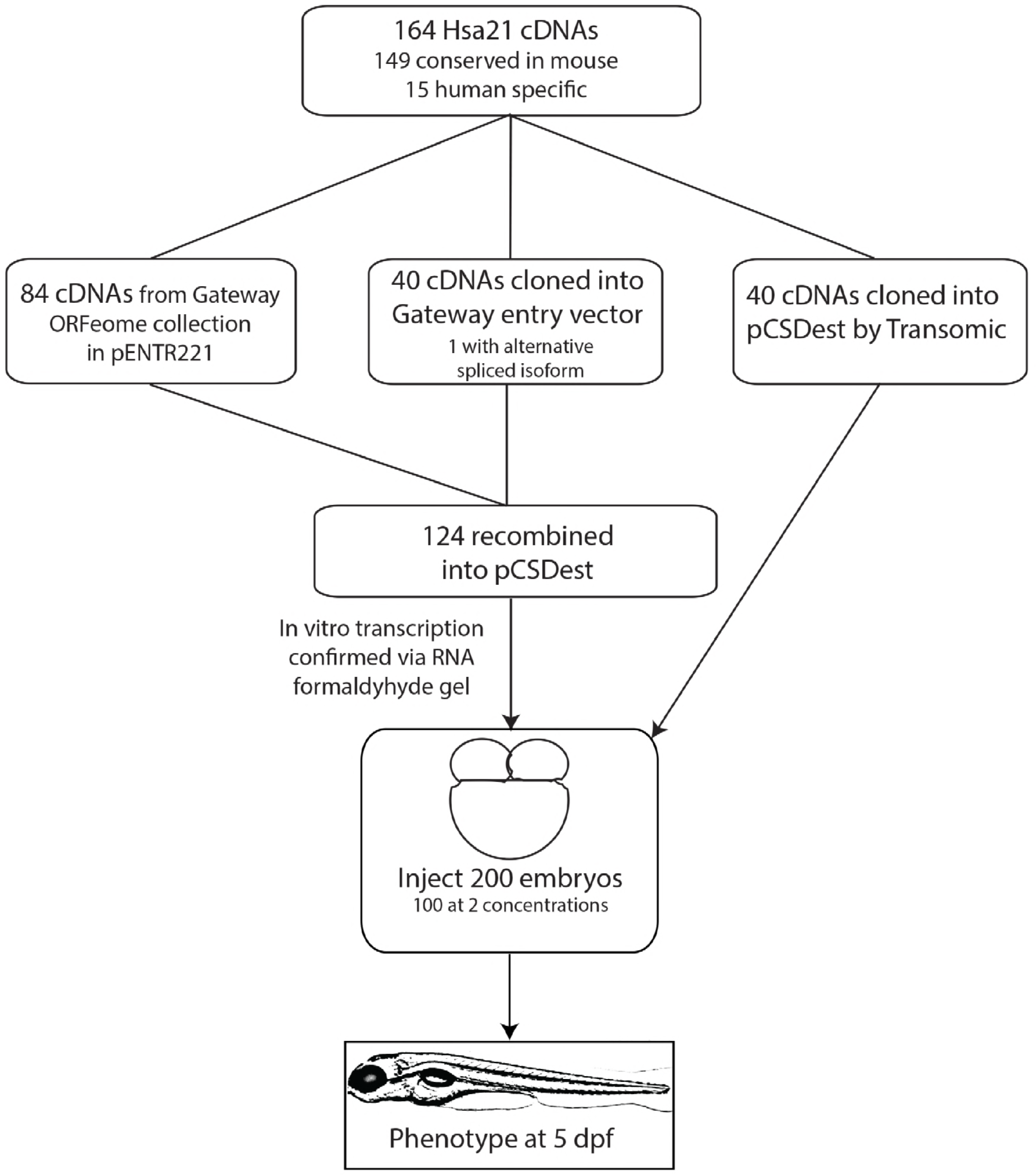
Flowchart showing steps of making the Hsa21 Gene Expression Clone-set and the screen of the set in zebrafish

Next we used Mouse Mine to interrogate the human gene list and compare it to homology data sets, HomoloGene and Panther, to identify orthologs and ohnologs between zebrafish and mouse (Supplemental Table 1) [23], [24]). In zebrafish, 125 of the 163 Hsa21 genes (77%) are conserved; 35 of those (28%) are represented by more than one zebrafish ortholog (ohnolog). The number of Hsa21 genes conserved with zebrafish is similar to published reports for all human genes (71%), but the proportion of Hsa21 genes with ohnologs is less than the 47% rate for all human genes [25].

### Zebrafish Screen

We synthesized mRNAs from the Hsa21 cDNA clones and injected each one into ~100 zebrafish embryos at the 1-2 cell stage at 10, 50 and/or 100 pg (<u>Supplemental Table 2</u>), ranges that have been used in similar studies previously [20]. More than 50,000 embryos were screened for the presence of gross morphological changes present at frequencies greater than controls, typically at five days post fertilization (dpf), but also earlier if mandated by the presence of clear pathologies. We focused primarily on three broad phenotypic classes: a) U-shaped somites and cyclopia, two phenotypes associated frequently but not exclusively with defects in the ciliome and Shh signaling pathway; b) craniofacial abnormalities and pigment differences, which may be related to aberrations affecting neural crest cells; and c) pericardial edema, which can have a number of causes including a structural heart defect (<u>Figure 2, Table 1</u>). We recorded the number of surviving embryos, the phenotype, and the percentage that were affected (<u>Table 2</u>).

**Figure 2:**
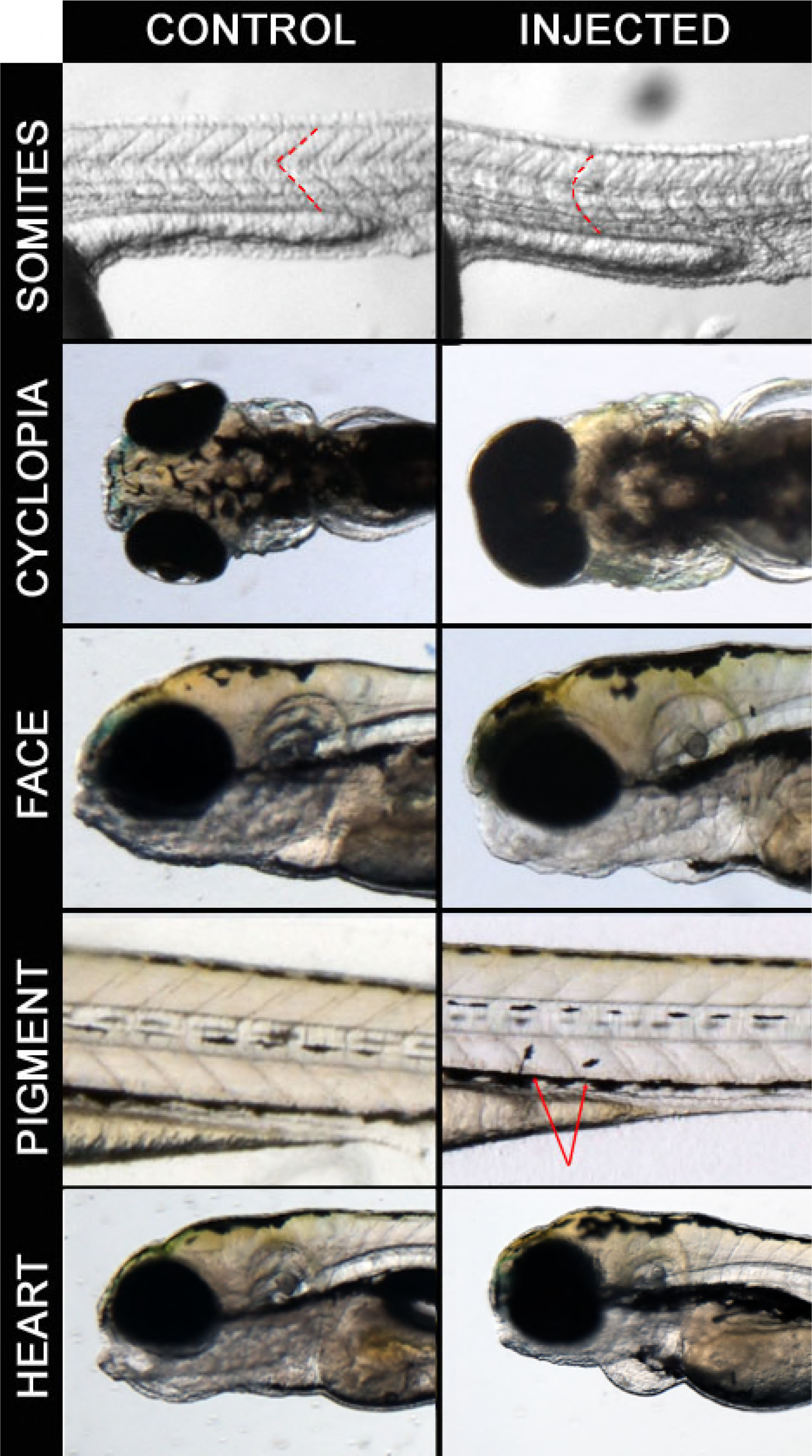
Examples of phenotypes observed in screen. Control embryos are on the left panel and injected embryos are on the right panel. Somites: *RWDD2B* 100pg injected embryos at 24 hpf with dashed lines to highlight somitic boundaries. Cyclopia: *C21ORF84* 100pg injected embryos at 5 dpf. Pigment cell migration: *CCT8* 100pg injected embryos at 4 dpf, arrows indicating melanocytes. Heart: *JAM2* 100pg injected embryos at 48 hpf.

**Table 1:** Twenty-four candidates from first pass of screen (see Supplemantal Table 3)

| **Phenotype category** | **Description** | **# Candidates** |
| --- | --- | --- |
| U-shaped Somites | Somites with characteristic U shape | 8 |
| Cyclopia | Single large eye | 6 |
| Craniofacial abnormalities | Small/missing mandible; skull abnormalities | 4 |
| Pigment abnormalities | Floating melanocytes; reduced pigment in eye | 3 |
| Heart | Pericardial edema | 1 |
| Other | Tail/fin abnormalities, embryo lethality | 2 |

**Table 2:** Final Candidate list of genes that produced a phenotype consistently (see also <u>Supplemental Table 2</u>).

| **Gene Symbol** | **Phenotype** | **Penetrance** | **Expression in mouse [31]** |
| --- | --- | --- | --- |
| *SOD1* | U-somites | 10-35% | Expressed ubiquitously at E10.5, strongly expressed in muscles at E14.5 |
| *RWDD2B* | U-somites | 15-31% | Expressed ubiquitously at E10.5 |
| *RRP1* | U-somites | 15-40% | Weakly expressed in somites at E10.5 [32] |
| *PCBP3* | U-somites | 8-13% | Expressed in brain and spinal cord at E14.5 |
| *YBEY* | U-somites | 12-19% | Expressed ubiquitously at E10.5 |
| *C21orf84* | U-somites/<br>Cyclopia | 9-23% U-somites<br>0-7% cyclopia | human specific [21] |
| *POFUT2* | Cyclopia | 0-4% | Strong in face and pharyngeal arches at E9.5 [32] |
| *CBR3* | Craniofacial | 7-10% | Expressed ubiquitously at E10.5, strong in cartilage at E14.5 |
| *CCT8* | Pigment | 15-40% | Expressed ubiquitously at E10.5 |
| *JAM2* | Pericardial edema | 20-60% | High expression in human heart [49] |
| *ERG* | Embryonic lethal | 64-93% | Expressed in the pharyngeal arches and limb buds at E10.5 [17] |

Of the 164 RNAs, 24 showed a phenotype after the initial screen (Supplemental Tables 2 and 3); the remaining 140 did not yield a significant number of affected embryos. Thus, again consistent with previous studies [20], our approach is not generally toxic to zebrafish embryos. Expression of 14 of these 24 genes gave a phenotype consistent with perturbation of the ciliome/Shh- pathway (eight with U-shaped somites and a partially overlapping set of six with cyclopia). An additional seven genes resulted in phenotypes that may be due to neural crest defects (four with craniofacial abnormalities and three with pigment differences). Finally, one gene induced pericardial edema, one resulted in dysmorphic fins, and one exhibited elevated lethality (<u>Table 1 and Supplemental Table 2</u>). Of the 24 first-round candidates, 19 are conserved in both mouse and zebrafish; four are conserved in between human and mouse but not zebrafish (*BACH1*, *Clic6*, *MAP3K7CL*, *SPATC1L*); and one gene (*LINC00313*) is human specific. Six of the 19 conserved Hsa21 genes have ohnologs in zebrafish (Supplemental Table 1).

Fresh mRNA was prepared from these first-round candidates and injections were repeated. Eleven candidates recapitulated the original phenotypes robustly, seven from the Shh group, two with neural crest related phenotypes, and the single genes resulting in pericardial edema and embryonic lethality, respectively (<u>Figure 2</u>, <u>Table 2</u>, <u>Supplemental Tables 2 and 3</u>). Of the 11 second-round candidates conserved in zebrafish, two have ohnologs in zebrafish (*JAM2* and *CBR3*).

### Phenotypic rescue via Morpholinos

The first indication that the phenotypes observed in repeated injections are due to the expression of the specific RNA and not to a general RNA effect is that there are at least 140 clones that produced no phenotype in this screen. Next, we selected randomly four of the candidate genes and performed a rescue experiment using morpholino (MO)-based knockdown. We designed translation-blocking MOs against the human copies of *SOD1*, *RWDD2B* or *CCT8* to target the ATG start site of the gene to suppress specifically the introduced human mRNA. We also designed MOs to knock down the endogenous zebrafish orthologs of *JAM2*, using a previously validated MO [26]. One hundred embryos were injected with 2ng MO and 100pg of RNA. For all four genes, injection of the RNA alone produced significantly higher penetrance than the uninjected controls, the MO alone, or the MO+RNA injected embryos (p<0.05 <u>Figure 3</u>). For *SOD1*, *RWDD2B*, and *CCT8* the MO+RNA was not significantly different from the controls. *JAM2* MO+RNA showed significantly lower penetrance of heart edema relative to RNA-injection alone [27]. The *JAM2* experiment suggests that this phenotype is a product of additive effects of the human cDNA with its fish orthologs/ohnologs.

**Figure 3:**
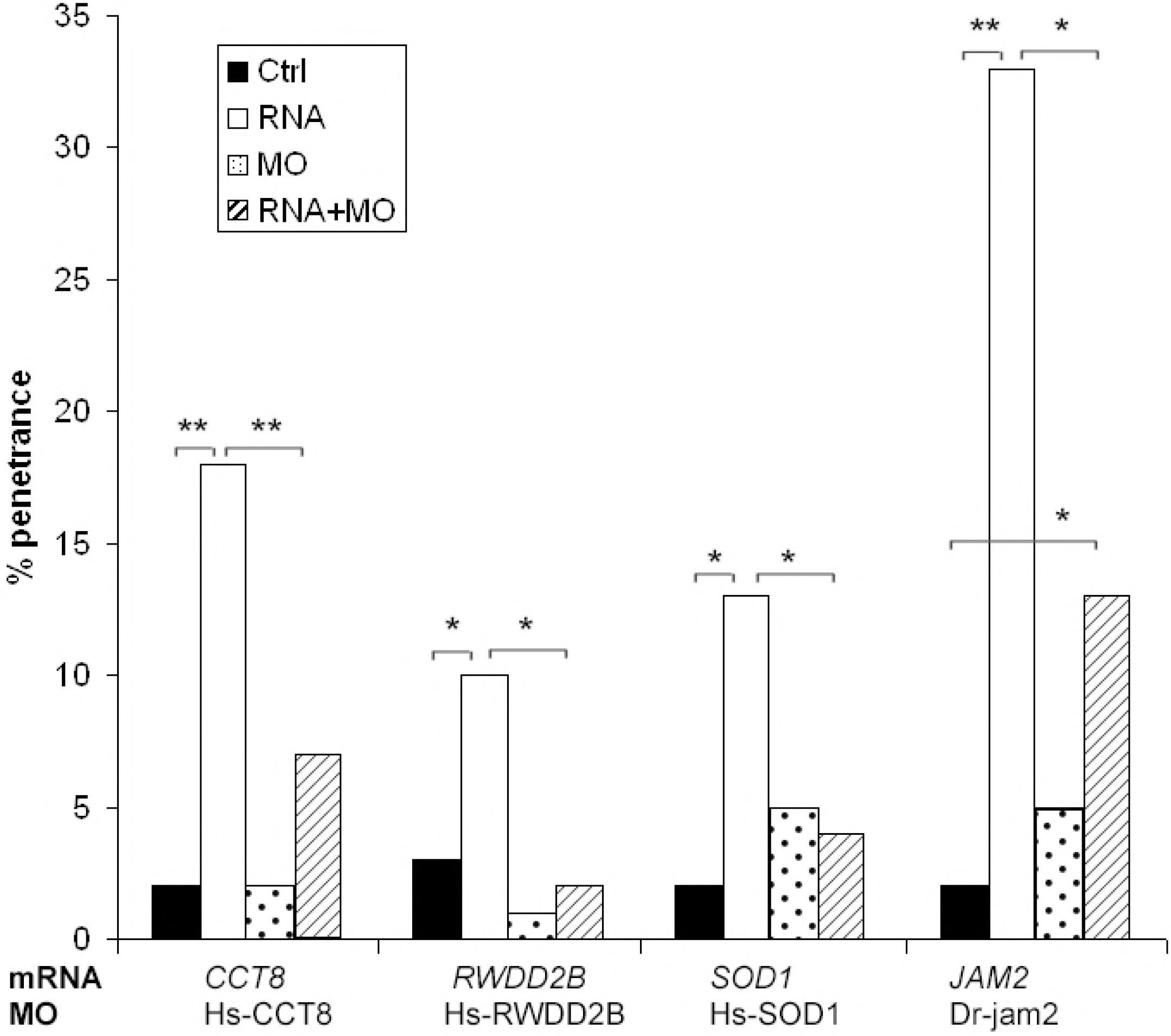
Candidate genes (*SOD1*, *RWDD2B*, *CCT8*, and *JAM2*) coinjected with translational blocking morpholino. Hs-*RWDD2B,* Hs-*CCT8* and Hs-*SOD1* MOs were targeted against the human mRNA, while *jam2* MOs were targeted against the zebrafish ortholog, DR-jam2. 100pg RNA was injected alone, 2 ng MO alone, or both were coinjected. * p<0.05, ** p<0.01. *JAM2* data adapted from [27].

### Dosage and Gene expression patterns

This screen was designed to identify Hsa21 genes with an effect on broad aspects of early embryonic development and not as a study of human dosage effects on fish (see Discussion). For those cDNAs producing phenotypes, penetrance was generally correlated with but not directly proportional to penetrance (<u>Supplemental Table 2</u>). In the case of *ERG*, however, the frequency of embryonic lethality was proportional to RNA dose. Approximately 1/3 of embryos (24/67) survived after injection with 10 pg ERG RNA, while 14% survived injection with 50 pg and 6.5% (24/358) were alive two days after injection of 100 pg. Retrospective examination also showed somewhat elevated mortality following injection of either of the additional two ETS family transcription factors in the Hsa21 clone set, *ETS2* and *GABPA* (<u>Supplemental Table 2</u>). *HMGN1* also showed a trend toward higher lethality.

Among the final candidates with repeatable phenotypes, only *ERG* showed increased lethality with increasing RNA concentrations. The 10 remaining candidates were injected at three or more concentrations ranging from 10pg to 200pg but no correlation between dosage and penetrance was observed (<u>Supplemental Figure 1</u>). The mRNA concentrations that we used are consistent with those commonly reported [20, 28]. We also chose two genes, *SOD1* and *RRP1*, to look for effects of low RNA doses. These genes were injected at 2pg and 5pg, and the penetrance of U-shaped somites was examined. Both genes showed low penetrance of the phenotype at these low doses, although each reproducibly produced a phenotype after injection of 50-100pg (<u>Supplemental Figure 2</u>).

Nine of the eleven candidates have zebrafish homologs, the exceptions being *C21ORF84* and *CBR3*. For eight of these genes, representative *in situ* hybridization data are available in ZFIN [29]. In most cases, the structure(s) affected by RNA injection is consistent with the *in situ* expression data. For example, *sod1, rrp1* and *ybey* are expressed in somites, *cct8* is expressed ubiquitously, *jam2* shows expression in the region of the developing heart and *erg* is expressed in the vasculature (Table 2). *pofut2* is expressed in the brain and eye [30]; knockdown of a *C. elegans* homolog of this gene produced a neuromuscular phenotype [15]. *pcbp3* is expressed in the retina and telencephalon beginning at the Prim 15 stage [29]. No information about *rwdd2b* expression was available in ZFIN. Ten of the eleven candidates have mouse orthologs whose expression has been examined at mid-gestation showing that the genes are expressed in the corresponding tissues during embryonic development in mouse (<u>Table 2, [31, 32]</u>). *C21ORF84* is a human specific lncRNA [21].

### Combinatorial injections

DS is a contiguous gene defect with effects on development that exceed those of individual genes. Examination of all pairwise interactions of 163 genes would require more than 13,000 pairs, posing a logistical challenge even in this system. We took an initial step toward understanding this process with respect to a small set of candidate genes. We first examined the subset of genes that produced possible Shh-related phenotypes in our screen. *SOD1*, *RWDD2B*, *YBEY*, *PCBP3*, and *RRP1* all produced U-shaped somites, *POFUT2* resulted in cyclopia and *C21ORF84*-injected embryos showed both phenotypes. We used *C21ORF84* as the reference gene to pair with the other six genes. For each set of pairwise injections, *C21ORF84* was injected alone at 100pg, the second gene was injected alone at 100pg, and both genes were injected together at 100pg each.

Injection of *C21ORF84* plus *YBEY*, *PCBP3* or *POFUT2* showed a significant increase in the frequency of embryos with U-shaped somites (Fig. 4, p<0.05 for each combination). For two genes, *SOD1* and *RWDD2B*, there was not a significant difference between the individual injections and the combinatorial injection. *RRP1* alone had a penetrance of 42%, the highest of all genes tested. This frequency was reduced significantly in embryos injected with *RRP1* and *C21ORF84* together.

**Figure 4:**
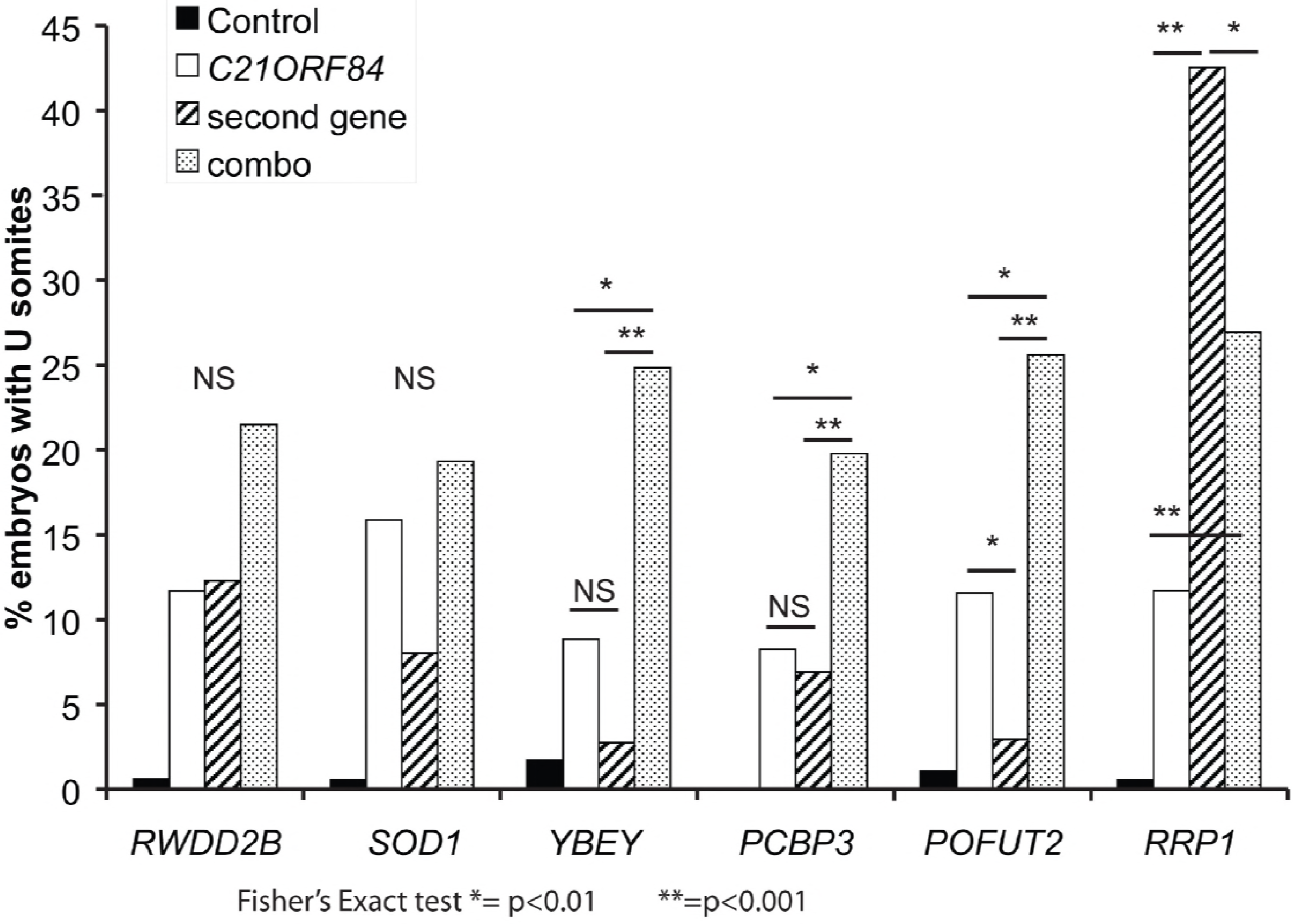
Pairwise combinatorial injections of Shh candidate genes. *C21ORF84* was coinjected with 6 other genes to look for synthetic effects. *C21ORF84* was injected individually at 100pg RNA, the other gene was injected individually at 100pg RNA and then the two were injected together, 100pg each, for a total of 200pg RNA.

We repeated the pairwise injections using a different reference gene from this group, *SOD1*. As in the previous experiment, *SOD1* was injected individually and in combination with one of the other six genes, and the embryos were examined for the presence of U-shaped somites. In contrast to *ORF84*, *SOD1* did not show an interaction with any of the other six genes (<u>Supplemental Figure 3</u>). Finally, combinatorial injections using *C21ORF84* were repeated with freshly prepared RNAs at a separate institution (by NAZ and CCL) as a means of independent replication. We observed a similar increase in affected embryos upon injection of *C21ORF84* with *YBEY* and *PCBP3* compared to each gene alone.

We also selected candidate gene sets for combinatorial injection based on reported roles in other systems and assessed them for possible combinatorial effects on developmental phenotypes. Several genes associated previously with heart anomalies in Down syndrome, *SH3BGR*, *DCSR6* and *ADAMTS1*, were co-injected (30pg each). This resulted in cyclopia in 3.6% and pericardial edema in a non-overlapping 3.6% of embryos, whereas no controls were observed to have either edema or cyclopia, a trend though not formally significant (Fisher’s exact test, p=0.056 for either cyclopia or edema). Several Hsa21 gene combinations have been implicated in the high frequency of congenital heart disease in DS [33]. However, neither injection of all three collagens together (*COL6A1*, *COL6A2*, *COL18A1*) nor coinjection of *DSCAM* and *SH3BGR* produced a significant frequency of heart or other gross anatomical defects.

## DISCUSSION

We have developed a study set of Hsa21 gene expression clones (available from AddGene, see Methods) and used it to conduct the first large-scale study of the effects of Hsa21 gene expression on early vertebrate embryogenesis. Previous analyses of Hsa21 gene expression in early development used *in situ* hybridization [31, 34] or microarrays to examine the localization, timing, and/or levels of Hsa21 gene up-regulation [2]. Here, we used a functional assay in zebrafish embryos to find candidate genes with effects on early development in a systematic, unbiased approach. This stage of development is difficult to study in mammalian models. Of the 163 genes (164 cDNAs) assayed, only eleven genes consistently showed an effect. These genes are implicated in ciliome/Shh signaling, neural crest cell (NCC) generation and differentiation, heart development, and/or embryonic lethality. *ERG* has recently been shown by the Knock-Out Mouse Phenotyping Project (KOMP2) to be required during mammalian embryogenesis; embryos homozygous for a null allele of *Erg* are lethal prior to E15.5 (www.mousephenotype.org accessed March 27, 2017 [35]). Several of the genes have little or no known function and none have been implicated previously in either Shh signaling or NCC generation and development. Notably, the *POFUT2* gene, was also found to have a neurobehavioral phenotype in a reciprocal loss-of-function screen in *C. elegans*, indicating that it may be a good candidate for further study [15].

There are several caveats to interpretation of this large scale screen. A negative result in the screen does not rule out a contribution of that gene to DS. First, some human genes could fail to produce an effect due to inefficient translation related to the fact that these are mammalian mRNAs that may not be properly processed in fish. Some human proteins may simply be hypo-or non-functional in zebrafish. Except for *SYNJ1*, we selected a single isoform for the cDNA clone set. Next, DS is a contiguous gene syndrome, with many interactions among the over-expressed genes and with disomic genes at different stages of development [11, 36]. Finally, it should be clear that *this is not a study of the effects of dosage imbalance* that occur in DS *per se*. Indeed, it would be difficult to define what dose of a human mRNA might result in human protein levels that, together with the orthologous fish proteins, replicate the *functional* stoichiometry of DS in a fish embryo. Genes with ohnologs that function in a complementary manner in fish represent an even more complicated situation when additional expression of (possibly) related but non-identical function is introduced via the human gene product. We did not observe a lower frequency of phenotypes among genes with ohnologs than in the Hsa21 gene set as a whole, but the numbers are very small. What we present is instead a screen for genes whose effects in this assay indicate candidates to pursue in far more labor intensive studies of early mammalian development. We have succeeded in one such application with *Jam2* [27].

The consistent occurrence of heart edema in zebrafish after injection of JAM2 RNA led us to examine this candidate in conjunction with increased penetrance of heart defects in trisomic mouse models. Ts65Dn “Down syndrome” mice are trisomic for about 104 genes orthologous to Hsa21 [37]. Breeding a null allele of the disomic gene, *Creld1*, onto Ts65Dn significantly increased penetrance of septal defects in the heart from 4% to 33% [38]. However, putting the same null allele on a related trisomy (Ts1Cje) that was trisomic for about 81 of the genes triplicated in Ts65Dn, had no impact on penetrance. *Jam2* was one of the 23 genes that are trisomic in Ts65Dn but not Ts1Cje, and one of 14 of these 23 that are expressed prenatally and/or in heart. Based on its effect on heart development in the zebrafish screen, we tested its role in mice by introducing a null allele of *Jam2* to produce Ts65Dn;*Creld1*+/−;*Jam2+/−* mice (returning *Jam2* to the normal two copies). Instead of the expected increase in penetrance in Ts65Dn;*Creld1*+/−, trisomy for all of the Ts65Dn genes except *Jam2* resulted in penetrance in these mice that was the same as Ts65Dn alone (i.e., 4%) [27]. Thus *Jam2* is a trisomic potentiator of the disomic modifier of heart disease penetrance, *Creld1*. This was the first demonstration of this type of genetic relationship in Down syndrome. It would not have been possible to pursue this relationship for all 14 candidate genes trisomic in Ts65Dn but not in Ts1Cje in mice.

In this screen, it was somewhat surprising that several Hsa21 genes that have been associated with robust phenotypes in mouse models of DS did not produce a phenotype. For example, *DSCAM*, a cell adhesion molecule that is involved in cell recognition [39], has been implicated in both heart and neurogenesis defects based on work done in mice as well as in *Drosophila* [40, 41] but did not produce a phenotype in our zebrafish screen. *DYRK1A*, a dual specificity kinase expressed during early neurogenesis that has been a target in pilot studies for treatment of cognitive deficits in DS [42, 43], also produced no phenotype. Many types of refined screens with greater sensitivity and specificity are possible, taking advantage of transgenically marked zebrafish lines to ask specific questions about development of specific structures. Furthermore, our identification of phenotypes associated with pathways previously identified as central to multiple manifestations of trisomy, such as Shh signaling [44–46], support the use of *D. rerio* for this type of large-scale systematic screen. This system represents a useful screening tool to identify individual candidate genes that may be significant drivers of DS phenotypes.

We observed that some Hsa21 genes that produce phenotypes on their own can interact in an additive manner, some have no apparent interaction and one pair had a compensatory interaction. Compensatory interaction implies that in some cases overexpression of one gene can balance the increased expression of another. To date, several genes of major affect have been associated with manifestations of Down syndrome in the mouse models. However, no single gene has been identified that is sufficient to produce completely a complex developmental phenotype of DS, consistent with the understanding of DS as a product of complex multi-gene interactions. Given the large number of possible gene-gene interactions on chr21 alone, the system described here provides a useful way to interrogate more complex interactions of non-contiguous genes from the earliest stages of development.

## METHODS

### Hsa21 Gene Expression Library preparation

cDNAs were selected from lists of conserved genes on Hsa21 [2, 21]. For 84 genes, plasmids containing the gene in the pENTR221 entry vector were obtained through the Invitrogen UltimateORF collection. Of the remaining genes, 49 were subcloned from a variety of vectors into one of the Invitrogen Gateway entry vectors (for complete list of original vectors and sources see <u>Supplemental Table 1</u>) and 40 genes were commercially cloned into the pCS2DEST vector. Entry vector clones were selected using kanamycin and then sequenced to confirm correct insertion of the gene. All genes in the entry vector were subcloned into the pCS2DEST vector (Addgene) using LR clonase as previously described by Katzen *et al.* 2007 [47]. Genes in the pCSDEST vector were selected by ampicillin. The entire pCS2+ vector clone set, named the Hsa21 Gene Expression Set, is available through Addgene (https://www.addgene.org/Roger_Reeves/). A limited set of HH pathway-related genes was used as a training set for the system including recognition of the U-shaped somite phenotype.

### Bioinformatics

Comparisons of human, mouse, and zebrafish orthologs and ohnologs was performed using MouseMine (www.mousemine.org, accessed October 1, 2017 [23]. Briefly, Hsa21 gene symbols for the 163 genes in this screen were uploaded as a list in MouseMine and interrogated against the mouse and zebrafish using the HomoloGene data set from NCBI and the PANTHER data set from MGI [24]. These lists were used to compile the ortholog and ohnolog lists in Supplemental Table 1.

### *In Vitro* Transcription of mRNA

Plasmids were transcribed *in vitro* using the mMessage mMachine SP6 kit (Ambion, Austin, TX). Plasmids were linearized and purified by precipitation. Transcribed sequence reactions were treated with DNAse1 and mRNA was purified with lithium chloride. mRNA quality and quantity were confirmed with a formaldehyde agarose gel and the Nanodrop8000, respectively.

### Zebrafish maintenance and injections

All procedures were approved by the Johns Hopkins University Animal Care and Use Committee, protocol no. FI15M197. Zebrafish were raised in the FINZ center at the Institute for Genetic Medicine (Johns Hopkins University) as described previously [48]. Zebrafish were maintained at 28°C. Male and female Tubingen zebrafish were placed in the same breeding tank in the morning and embryos were collected 30 minutes later. One hundred embryos were then injected at the 1-4 cell blastula stage using a Zeiss Stemi 2000 microscope and PV820 Pneumatic picopump injector. All genes were injected at 50pg or 100pg and most were injected a second time at a different dose (Supplemental. Table 2). All of those producing a phenotype were re-injected at 100pg. Embryos were raised to 5 days post fertilization and then phenotyped using a Nikon SMZ1500 microscope and imaged with NIS Elements Imaging Software. After imaging, embryos were fixed in 4% PFA overnight then transferred to 100% Methanol for storage at −20°C. For low dosage experiments, *SOD1* and *RRP1* were injected at 2pg, 5pg and 10pg and examined at 5 dpf.

### Morpholino Rescue

Translation-blocking antisense morpholinos (MO) were designed against the human sequence for the genes *SOD1*, *RWDD2B*, and *CCT8*, designed to bind to the ATG start codon of the mRNA using Gene Tools (Philomath, OR): Hs-*SOD1* 5’-GCACGCACACGGCCTTCGTCGCCAT-3’; Hs-*RWDD2B* 5’-GCTGCATGGACAGCTCAATTTTCAT-3’; and Hs-*CCT8* 5’-GAGCCTTGGGAACGTGAAGCGCCAT-3’. The MOs were checked using BLAST for sequence specificity to the human homolog and insure that they were unique in the either the human or zebrafish genomes. For each gene, 100 embryos were injected with 2ng MO, 100 embryos were injected with 100pg of mRNA and 100 embryos were injected with both 2ng MO and 100pg mRNA; 100 uninjected embryos were used as a control. Embryos were examined at 5 dpf for Hs-*SOD1* and Hs-*RWDD2B*, 4 dpf for Hs-*CCT8*, and 24 hrs. post-fertilization (hpf) for Dr-*JAM2*.

### Combinatorial Injections

RNA from *C21ORF84* was coinjected with RNA from the following genes: *SOD1*, *RWDD2B*, *RRP1*, *PCBP3*, *POFUT2*, and *YBEY*. Each gene was injected individually at 100pg into 100 embryos, and then coinjected at 100pg of each RNA (200pg RNA). Embryos were phenotyped at 24 hpf for the presence of U shaped somites and cyclopia. Each coinjection was performed twice. The entire experiment was carried out independently at a different institution (NAZ and CCL) injecting 200 pg mRNA into 50-100 embryos of the Tubingen line using the pairs listed and phenotyping at 24 hpf.

The combinatorial strategy was repeated using *SOD1* as the reference gene and coinjected with the same genes listed above. In this case, 50pg of each RNA was injected individually into 100 embryos each, and then 50pg each of both RNAs were coinjected into 100 embryos, with 100 control embryos. Embryos were examined at 24hpf for the presence of U-shaped somites.

### Statistical Tests

For all injections, penetrance differences were examined using a Fisher’s Exact test with p<0.05 required for significance.

## Acknowledgments

We thank Valerie DeLeon, Deborah Andrew, and Steven Leach for their advice and support. This work is submitted in partial fulfillment of the requirements for a Ph.D. (SE).

## Author Contributions

SE, RR, NK and AM conceived and designed the experiments. SE, NAZ, CCL, DK, JT and JL performed the experiments. SE, NAZ, RR and AM analyzed the data. SE and RR wrote the manuscript, which was reviewed and revised by all authors.

---

**Supplemental Figure 1:**
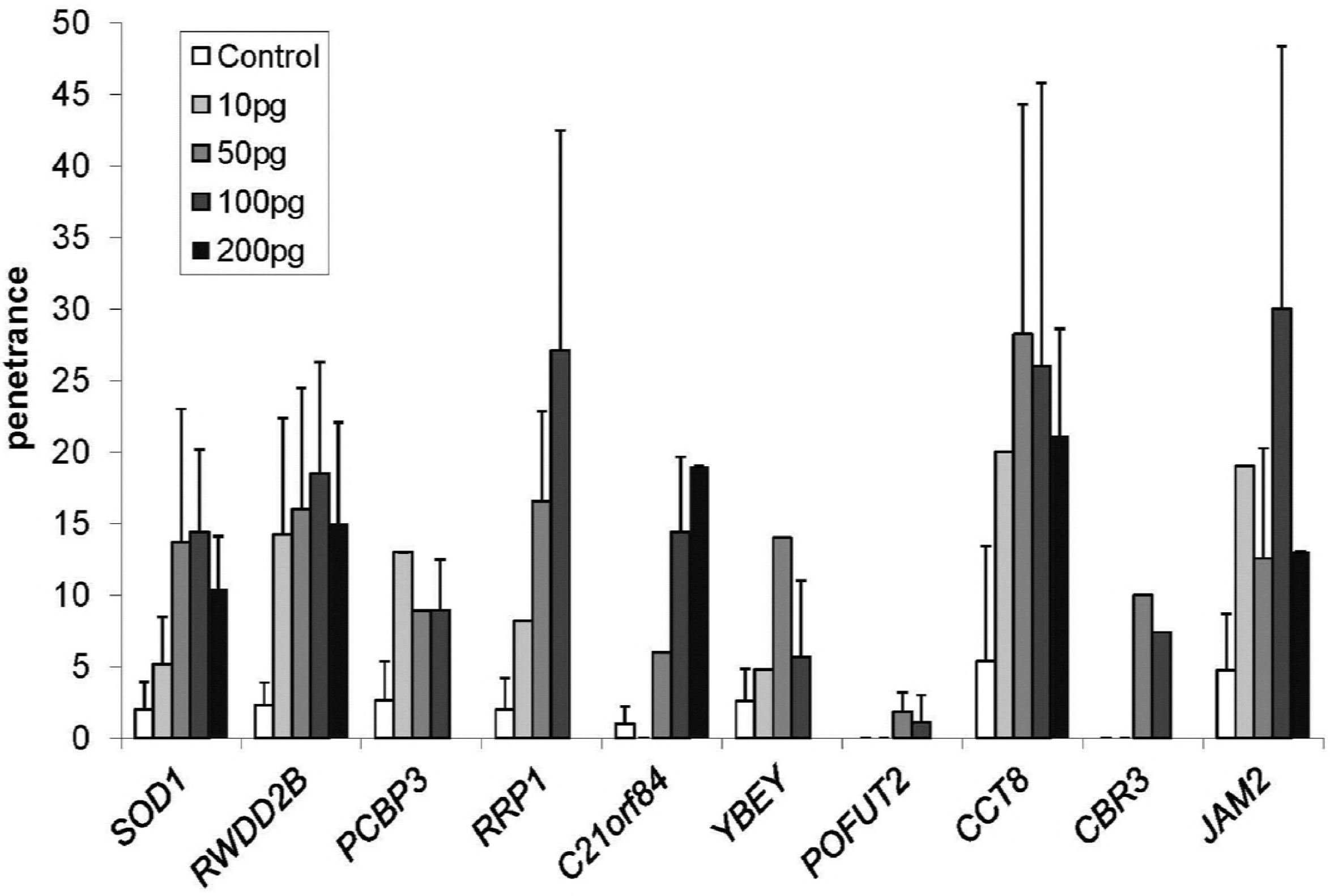
Human mRNAs did not show a dose response at the concentrations used. Phenotype penetrance is shown for the ten candidate genes at concentrations from 10-200 pg/embryo.

**Supplemental Figure 2:**
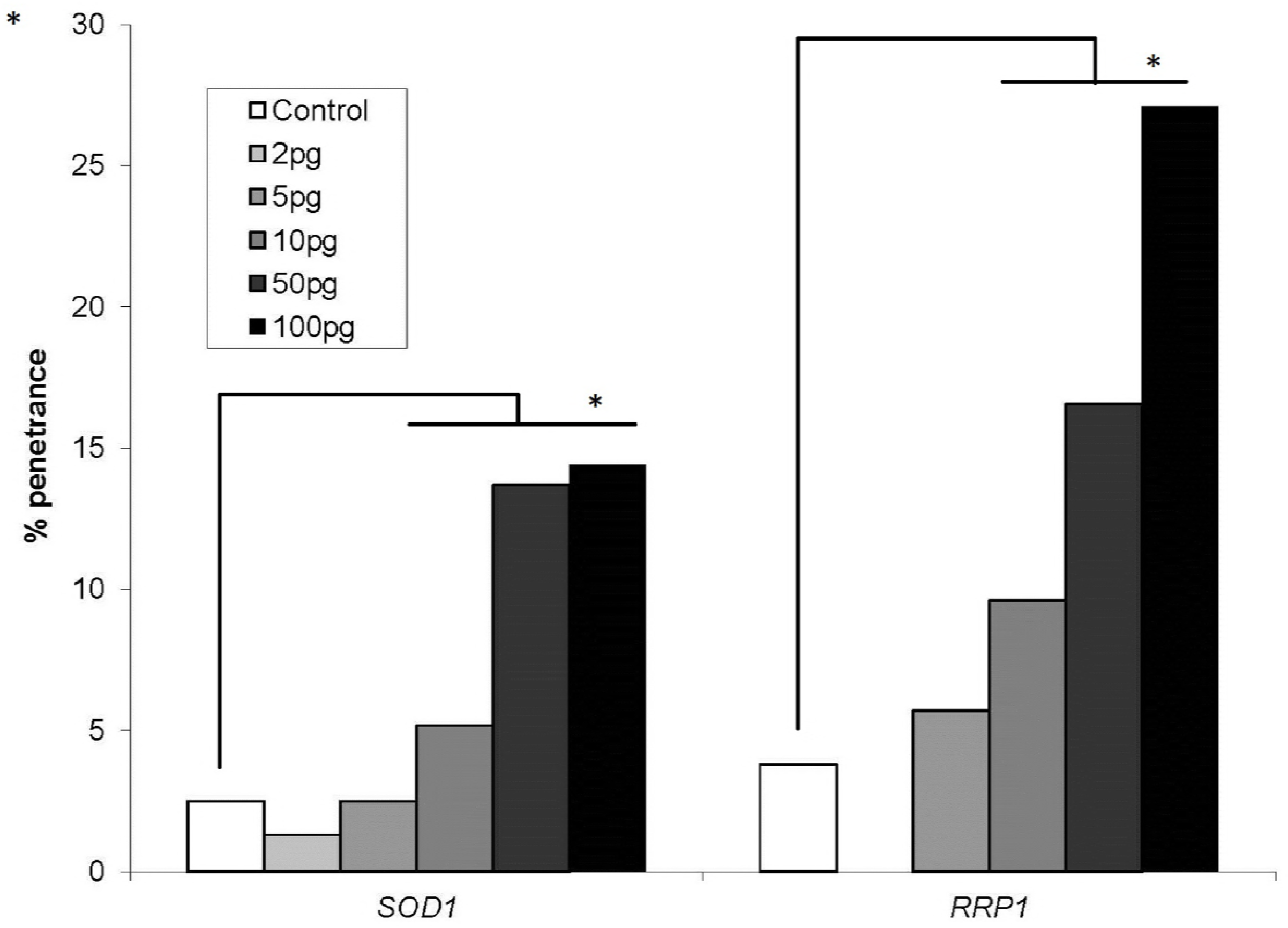
SOD1 and RRP1 were examined for penetrance of the phenotype at very low RNA doses but did not show significant penetrance. Uninjected embryos for controls. * p<0.01

**Supplemental Figure 3:**
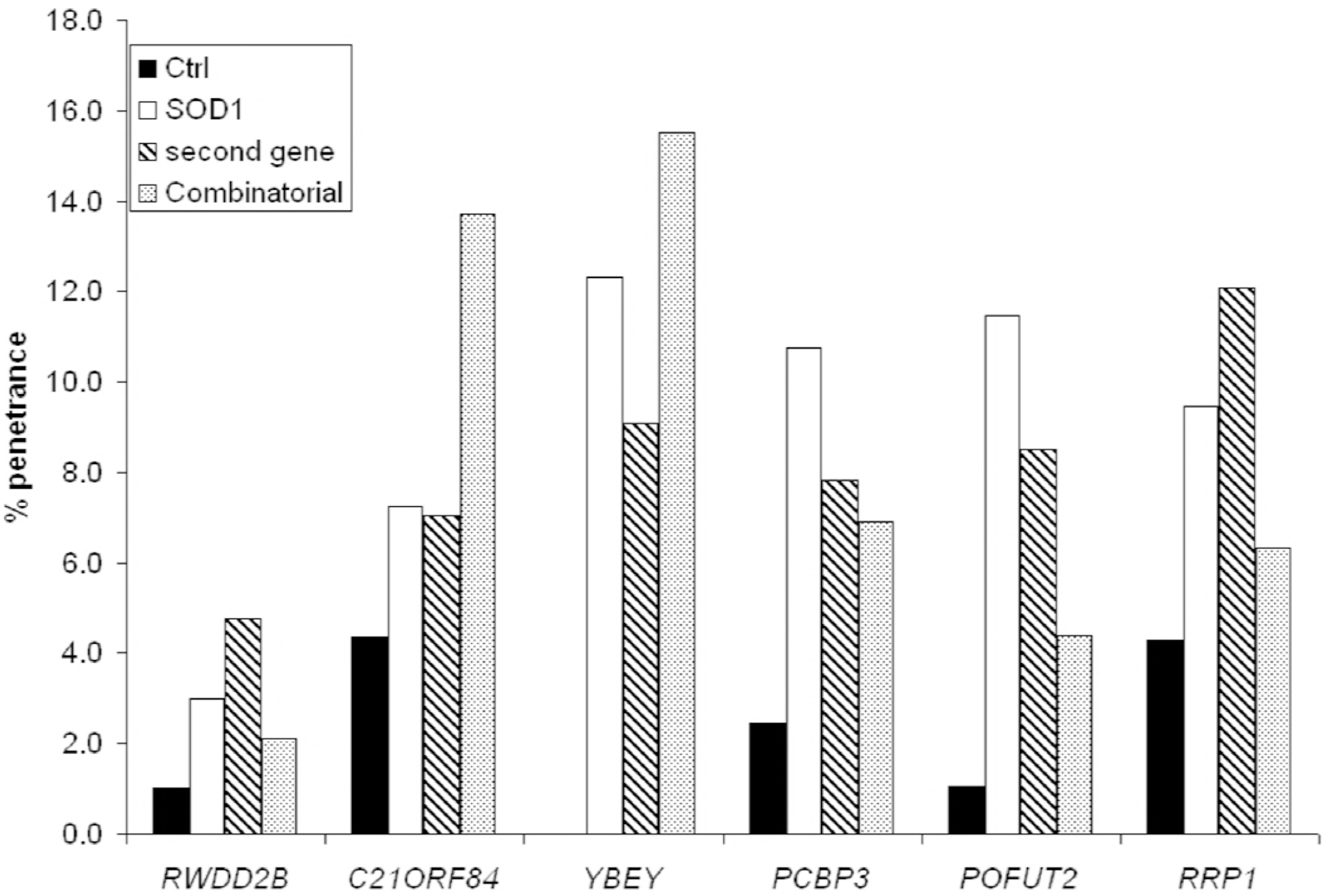
Pairwise injections of Shh related genes using *SOD1* as the reference gene. *SOD1* was coinjected with 6 other genes to look for synthetic effects. *SOD1* was injected individually at 50pg RNA, the other gene was injected individually at 100pg RNA and then the two were injected together, 50pg each, for a total of 100pg RNA.

